# Predicting clone genotypes from tumor bulk sequencing of multiple samples

**DOI:** 10.1101/341180

**Authors:** Sayaka Miura, Karen Gomez, Oscar Murillo, Louise A Huuki, Tracy Vu, Tiffany Buturla, Sudhir Kumar

## Abstract

**Motivation:** Analyses of data generated from bulk sequencing of tumors have revealed extensive genomic heterogeneity within patients. Many computational methods have been developed to enable the inference of genotypes of tumor cell populations (clones) from bulk sequencing data. However, the relative and absolute accuracy of available computational methods in estimating clone counts and clone genotypes is not yet known.

**Results:** We have assessed the performance of nine methods, including eight previously-published and one new method (CloneFinder), by analyzing computer simulated datasets. CloneFinder, LICHeE, CITUP, and cloneHD inferred clone genotypes with low error (<5% per clone) for a majority of datasets in which the tumor samples contained evolutionarily-related clones. Computational methods did not perform well for datasets in which tumor samples contained mixtures of clones from different clonal lineages. Generally, the number of clones was underestimated by cloneHD and overestimated by Phy-loWGS, and BayClone2, Canopy, and Clomial required prior information regarding the number of clones. AncesTree and Canopy did not produce results for a large number of datasets.

**Conclusions:** Deconvolution of clone genotypes from single nucleotide variant (SNV) frequency differences among tumor samples remains challenging, so there is a need to develop more accurate computational methods and robust software for clone genotype inference.

**Availability and Implementation:** CloneFinder is implemented in Python and is available from https://github.com/gstecher/CloneFinderAPI.

**Supplementary information:** Supplementary data are available at *Bioinformatics* online

## 1 Introduction

Next generation sequencing has revealed that genetically distinct tumor cell populations (clones) originate from pre-existing clones and change in frequency over time (de Bruin et al., 2014; Gerlinger et al., 2012; Nik-Zainal et al., 2012; Schuh et al., 2012). While cells may become cancerous through many types of micro- and macro-mutations, many studies currently analyze single nucleotide variants (SNVs) (Cancer Genome Atlas Research et al., 2017; Helleday et al., 2014; Horne et al., 2014). Ideally, researchers would generate a comprehensive catalogue of clone genotypes by individually sequencing all of the cells in a given tumor sample. This would enable the inference of the evolutionary history of clones, identification of early and late occurring driver mutations, and characterization of inter- and intra-tumor heterogeneity (Gundem et al., 2015; Hong et al., 2015; Macintyre et al., 2017; Turajlic et al., 2015). However, single cell sequencing remains expensive and technically challenging (Gawad et al., 2016; Navin, 2015; Sun et al., 2015). Consequently, tumor bulk sequencing is currently the fastest and most cost-effective means to assess the genetic heterogeneity of tumor samples (de Bruin et al., 2014; Nik-Zainal et al., 2012; Ojha et al., 2015; Schuh et al., 2012). Bulk sequencing produces SNV frequencies for the whole population of cells in each tumor sample. While phylogenies of tumor samples are commonly inferred using these data, these phylogenies are not identical to clone phylogenies because multiple clones are often present within each sample (Alves et al., 2017). Thus, the use of tumor phylogenies is not always appropriate for making biological inferences about the origin of clones and their evolutionary relationships.

Many computational methods predict clone genotypes from SNV frequency data produced by bulk sequencing of multiple tumor samples (Beerenwinkel et al., 2014; Vandin, 2017). The original descriptions of these computational methods provide an initial evaluation of their performance in recovering the relative temporal order and clustering of mutations (Deshwar et al., 2015; El-Kebir et al., 2015; Farahani et al., 2017; Malikic et al., 2015; Popic et al., 2015), the number of mutation clusters (Deshwar et al., 2015), and the topology of the clone phylogeny (Malikic et al., 2015). The clone genotype errors are also sometimes reported (Sengupta et al., 2015; Zare et al., 2014).

However, the performance of these methods has not been tested using uniform datasets and metrics for evaluation, such as the accuracy of the number of clones inferred and the quality of clone genotypes. Therefore, we have evaluated eight existing methods (Table 1) that predict clone genotypes based on the analysis of SNV frequencies (or read counts) produced by bulk population sequencing of tumor samples. All of these methods produce clone genotypes directly from the SNV frequency data. In addition, we developed and tested a method (CloneFinder) that employs principles of molecular phylogenetics to infer clone genotypes by analyzing data from bulk sequencing of multiple tumor samples (see Methods).

**Table 1.**
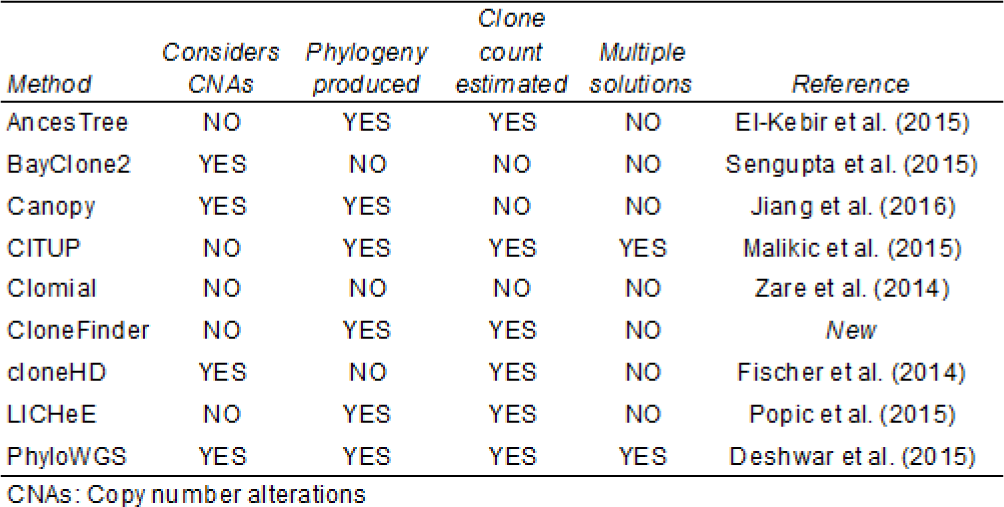
Nine clone prediction methods tested

Computer simulation and analysis of data is a useful tool for evaluating the performance of statistical methods in cancer genomics (Deshwar et al., 2015; El-Kebir et al., 2015; Farahani et al., 2017; Malikic et al., 2015; Popic et al., 2015). For simulated datasets, the correct numbers of clones and each clone genotype are known, enabling direct evaluation of the performance of computational methods. We also analyzed two empirical datasets (McPherson et al., 2016; Stachler et al., 2015) in order to test if the nine methods in Table 1 can produce the same inferences that were reported in the original empirical studies.

In the following, we present methods used for generating the test datasets, a description of the CloneFinder method, and the assumptions and parameters used while applying each of the nine computational methods. This is followed by definitions of the performance measures used to evaluate methods. We then present results from our analyses and compare these results with those reported in previous studies.

## 2 Methods

### 2.1 New method evaluated (CloneFinder)

We explain the CloneFinder method by using a G7 dataset consisting of seven clones and seven samples (Fig. 1a and 1b), which was used to generate observed SNV frequencies, V, in the tumor samples (Fig. 2a). V is a two-dimensional matrix in which the number of rows is equal to the number of tumor samples (i’s) and the number of columns is equal to the number of SNVs (j’s). Using this matrix, CloneFinder first constructs a matrix I (*M*) in which the presence of a SNV (V_ij_ > 0 observed SNV frequency) is represented by a 1, and the absence is assigned a 0 (V_ij_ = 0 observed SNV frequency). This *M* is the initial collection of candidate clone genotypes (Fig. 2b). Duplicate rows in *M* are then eliminated to keep one of the rows, which would happen when the same set of SNVs are present in some tumor samples. This was observed for samples T3 and T4 and samples T1 and T5 in our example dataset, so two of the rows in *M* were eliminated (Fig. 2c). Therefore, the number of initial clone genotypes (k’s) is five.

**Figure 1:**
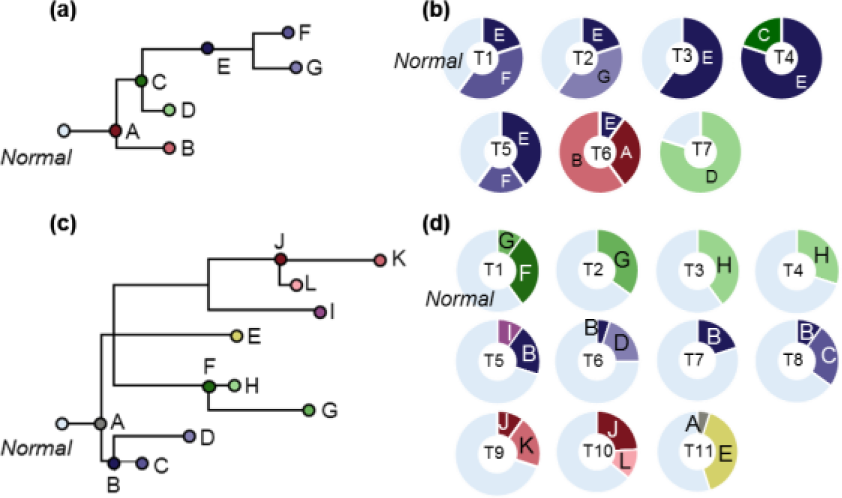
Clonal history and frequencies used as models generating G7 and G12 datasets. (**a**) A 7-clone phylogeny derived from EV005 tree in Gerlinger et al. (2014). (**b**) Clone frequencies in seven tumor samples (T1-T7) that were used to generate G7 datasets. (**c**) A 12-clone phylogeny derived from RK26 tree in Gerlinger et al. (2014). (**d**) Clone frequencies in eleven tumor samples (T1-T1 1) that were used to generate G12 datasets. All clones descend from the root clone A in each tree.

CloneFinder then tests whether the constituent clone genotypes are identical to tumor genotypes. If a sample contains only one clone genotype along with normal cells, then the genotype of the constituent clone will be identical to the tumor genotype. For this test, CloneFinder defines ½*f* × *M* = V, where *f* is a two-dimensional matrix of estimated clone frequencies of the tumor samples. This equation applies to cases where the variants are free of copy number alterations (CNAs). Otherwise, one may use estimated cancer cell fraction (CCF) for each variant site (Landau et al., 2013) as an input. In this case, *f* × *M* = V, where V_ij_’s are CCFs.

CloneFinder carries out regression analysis under the Karush-Kuhn-Tucker conditions (Kuhn, 1951) for non-negative least squares to generate estimates of *f*_ik._ If the difference between the observed (V_ij_) and the predicted (*v*_ij_) SNV frequencies is (nearly) zero (∂_ij_ = *v*_ij_ – V_ij_ ~ 0) for all the SNVs in all the tumor samples, then the binary vectors can be immediately translated into clone genotypes (Fig. 2d). In our example, ∂_3j_ = 0 for all SNVs in sample T3, which is reasonable because T3 contained clone E and normal cells. The same is true for sample T7 (clone D) (Fig. 2j).

**Figure 2:**
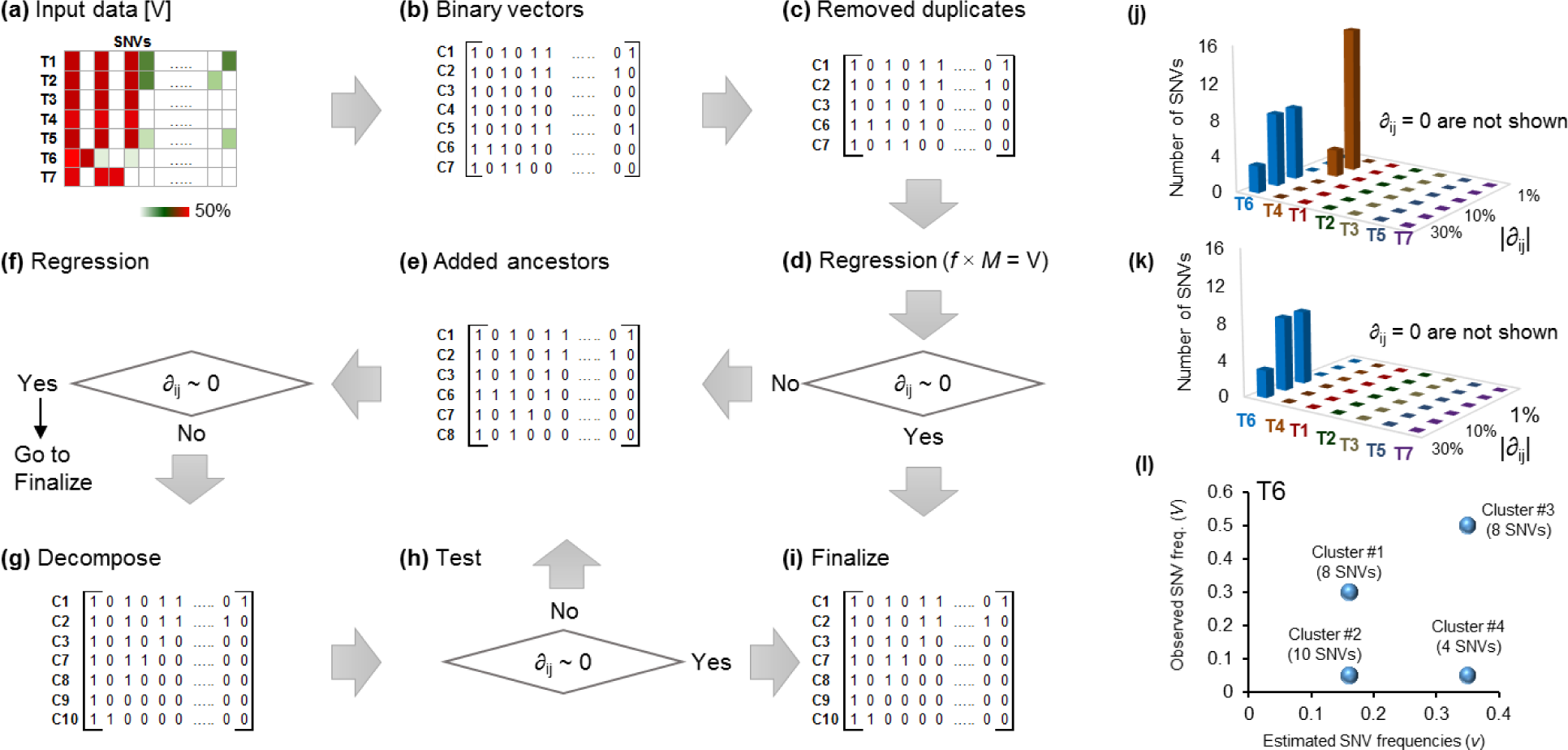
An overview of the CloneFinder method. (**a**) Input data of CloneFinder is observed SNV frequencies. A sample of observed variant frequencies of seven samples (T1–T7) is shown. **b**) Input data is transformed into the binary representation of presence and absence of mutations, which are the initial candidate clone sequences (genotypes). (**c**) Redundant candidate clone genotype are removed (C4 and C5). Also, regression analysis is used to eliminate clone genotypes present at zero or very low frequency in all samples. No clones were removed in this example. (**d**) Regression analysis is used to compute ∂. (**e**) Expanded list of candidate clones after the addition of ancestral and root clones. (**f**) Regression analysis. (**g**) Expanded list of candidate clones after the addition of decomposed clones. (**h**) Regression analysis. (**i**) Clone genotypes are finalized by removing SNVs affected by backward or parallel mutations. (**j** and **k**) SNVs showing differences between the observed and predicted frequencies (∂) in tumors for the candidate clone genotypes are in the panel **c** and **e**, respectively. The addition of ancestral clones resolves the discrepancy between the observed and predicted frequencies of SNVs in T4 in panel **k**. (**l**) Clusters of SNVs show different observed and predicted frequencies of SNVs in T6, and SNV clusters are combined to produce candidate clone genotypes in panel **g**. The number of SNVs within a cluster is shown in parenthesis. In panel **a**, we recommend pre-processing of input data to remove low-quality SNVs, e.g., using a Bayesian inference model to compute the posterior probability of the presence of variant (Reiter et al., 2017). Also, SNVs that are affected by copy number alterations (CNAs) should not be included when using the CloneFinder method.

This analysis also revealed that the difference between the estimated and observed SNV frequencies was zero for samples T1, T2, and T5, i.e., ∂_1j_ = 0, ∂_2j_ = 0, and ∂_5j_ = 0, respectively. However, these samples contained two different clones. They were discovered because one of their clone genotypes is also found in another sample, and the tumor-specific clone in each case has a genotype identical with the tumor genotype, e.g., T2 contains clones C2 and C3. In contrast, the regression analysis leads to a nonzero ∂_4j_ and ∂_6j,_ for samples T4 and T6, respectively (Fig. 2j). CloneFinder hypothesizes that these non-zero ∂_ij_’s are caused by the presence of ancestral clone genotypes that have persisted either because they have not been replaced by descendant clones or because they have adaptive significance.

CloneFinder uses the transpose of *M* for phylogenetic clustering and generation of ancestral states following the maximum parsimony principles (Nei and Kumar, 2000). Because the numbers of sequence differences and homoplasic sites are both expected to be small, the use of the parsimony principle is appropriate. We found that a Bayesian approach to inferring phylogeny and ancestral states (Yang et al., 1995) produces similar results (not shown).

In the current example, the maximum parsimony tree contained two internal nodes, which yielded two candidate ancestral clones with binary genotypes different from those already present in *M*. Their addition expands *M* by two. CloneFinder re-estimates clone frequencies (*f*) by linear regression, revealing that one of the clones in *M* does not occur at a significant frequency in any of the tumor samples. This clone is removed from *M* and the regression analysis repeated to produce ∂_ij_’s (Fig. 2e and 2f). The estimated SNV frequencies were identical to the observed SNV frequencies for T4 (Fig. 2k). However, many SNVs still show a large difference in T6. This suggests that some candidate clones in *M* are genotype hybrids, which need to be decomposed into constituent clone genotypes.

To decompose hybrid genotypes, CloneFinder first groups SNVs that have identical estimated SNV frequencies. In the case of T6, *v*_6j_ is 0.16, 0.35, or 0 for all the SNVs (Fig. 2l). CloneFinder pools SNVs with *v*_6j_ > 0 into two groups (*G*_0.16_ and *G*_0.35_) and tests if the SNVs within a group show significantly different V_6j_ (*t*-test; Fig. 2l). For example, SNVs in *G*_0.35_ show very different V_6j_’s (0.05 or 0.50), which results in the formation of subgroups consisting of 4 and 8 SNVs, respectively (Fig. 2l). In our example dataset, there were no read count errors, so the observed frequencies within a cluster are identical among SNVs. In the real world data, the observed frequencies will generally be different even within a correct cluster of SNVs. Nevertheless, we expect that observed frequencies of SNVs from the same cluster will be more similar to each other than to SNV frequencies in other clusters.

CloneFinder combines SNV clusters to produce new candidate clone genotypes. For example, one of the new candidate clone genotype will have SNVs from two clusters (#1 and #2 in Fig. 2l). CloneFinder generates all possible combinations of SNV clusters and finds the most likely combinations (correct clones) by using regression to estimate clone frequencies. CloneFinder expects that incorrect candidate clones will have effectively zero (i.e., low) clone frequency across all tumor samples, so they can be detected and removed. Only correct candidate clone genotypes were retained for the example dataset (Fig. 2g). Now, ∂_6j_ becomes zero, so the CloneFinder analysis ends with seven clones, all of which have genotypes identical to the true clone genotypes (Fig. 2h).

However, in the real world datasets, SNV frequencies produced by bulk sequencing will not be ascertained perfectly. So, CloneFinder applies the decomposition step described above only when at least one tumor sample shows a large ∂_ij_ and significant SNV clusters are detected by regression analyses. When this condition is met, CloneFinder continues to add candidate intermediate clones and refine clone genotypes iteratively until no additional clones are discovered. CloneFinder also stops if the new *M* increases the number of backward and parallel mutations in the clone phylogeny, which is an indication that spurious new clones are being added. In fact, many clone prediction methods do not allow backward and parallel mutations in their inferred clone phylogenies, e.g., (Malikic et al., 2015).

To finalize clone genotypes, CloneFinder applies a filter to SNVs that are predicted to be present on two independent clones within a sample, where it removes the SNV from the more recently emerged clone if the user desires (Fig. 2i). Also, CloneFinder has an option to filter backward / multiple mutations from the final clone sequences (i.e., wild-type bases are assigned at these positions).

### 2.2 Generation of datasets by computer simulation

#### The Gerlinger collection

(G7 and G12 datasets): We simulated two data collections using the evolutionary trees in Gerlinger et al. (2014) as model clone phylogenies (Fig. 1a and c). Up to 10 mutations were randomly assigned to branches in the clone phylogeny, which resulted in clone genotypes containing 34 – 88 SNVs. Tumor samples were populated with tip (e.g., B, D, F and G in Fig. 1a), ancestral (e.g., C and E in Fig. 1a), and root (e.g., A in Fig. 1a) clones in varying proportions drawn randomly (10% to 80%; Fig. 1b and d). One hundred simulation replicates each were generated using a 7-clone phylogeny (G7 datasets, Fig. 1a) and a 12-clone phylogeny (G12 datasets, Fig. 1c). The G7 dataset contained seven tumor samples and the G12 dataset contained 11.

#### The El-Kebir collection

(E10 datasets): We obtained 90 simulated datasets from El-Kebir et al. (2015), who generated random phylogenies containing 10 clones for each replicate. The simulated datasets contained 100 SNVs. Each dataset contained 4 – 6 tumor samples, and each tumor sample was populated with up to four distinct clone genotypes that were randomly chosen from the clone phylogeny. Since some clones may not be assigned to any tumor samples, the total number of clones in a dataset varied from 6 to 10. A total of 90 clone phylogenies were simulated, a sample of which is shown in Fig. 3.

**Figure 3:**
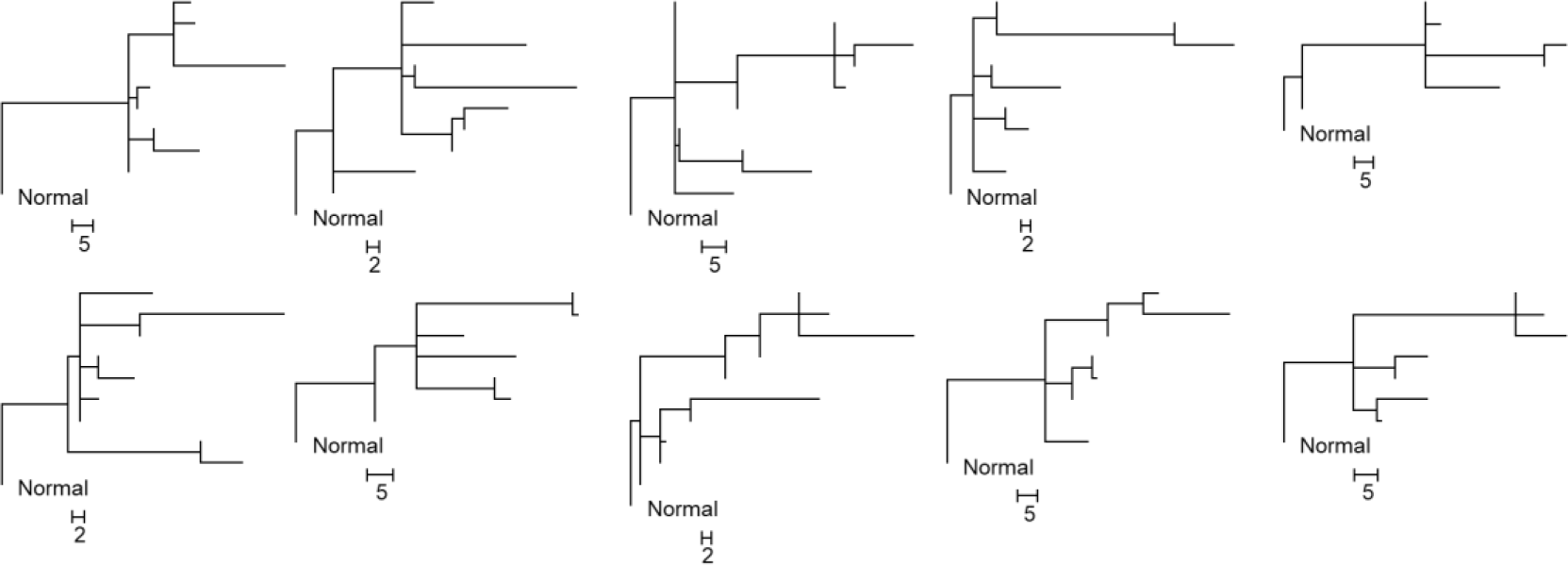
Ten clone phylogenies used for generating P10 datasets. Each dataset contained six tumor samples (six to 10 clones). Scale bars show the number of mutations along a branch.

#### The Popic collection

(P10 datasets). We assembled another 30 datasets from the El-Kebir et al. (2015) clone collections. Each dataset consisted of six tumor samples that were populated based on 10 different clone phylogenies (3 datasets each; Fig. 3). We used an approach similar to Popic et al. (2015)’ s “ localized sampling process” that is modeled after patterns seen in biopsies from spatially distinct sites within a tumor and/or among metastatic tumors. Their scheme populates tumor samples with closely-related clones based on the simulated clone phylogenies. In our simulation, we populated each tumor sample with one tip clone and its ancestral clones. We used clone frequencies from El-Kebir et al. (2015)’s datasets and assigned the largest clone frequencies to the tip clones.

#### Generation of read counts for G7, G12, and P10 datasets

The expected total read count for a SNV position was 100 in each tumor sample. Assuming that all mutations were heterozygous, we computed expected SNV frequencies. To generate SNV read counts with error (noise), we randomly sampled from a Binomial distribution with 100 reads (trials) and a given expected SNV frequency (*v*), i.e., Binomial (100, *v*). To introduce noise in the wild type read count, we randomly drew an observed total read count from a Poisson distribution with a mean of 100, and then subtracted the variant read count from it (see also Supplementary Note). SNVs were simulated without any genomic location considerations and every SNV was assumed to have been sequenced in a different read. We did not simulate CNAs because only four of the nine clone prediction methods are able to analyze SNV frequencies affected by CNAs (Table 1).

### 2.3 Accuracy measurements

We mapped each inferred clone genotype to its most similar true clone genotype in a two-step process. First, each inferred clone genotype was compared to all the true clone genotypes, and the two clones with the smallest difference were paired. This process was repeated using the remaining inferred clone genotypes and the remaining true clone genotypes. In this way, each inferred clone was matched to the most similar true clone. When the number of inferred clones was greater than the number of true clones, we first paired all true clones with the most similar inferred clones, and then the remaining inferred clones were paired with the most similar true clone genotype.

We counted the number of SNV assignment differences between the inferred and the true clone genotypes, which we refer to as the clone genotype error (*G*_E_). *G*_E_ varied from 0 to the number of SNVs in the dataset. %*G*_E_ is equal to *G*_E_ divided by the number of SNVs in the dataset. Multiple equivalent clone sets (multiple solutions produced for a dataset) were pooled together and all the distinct clone genotypes were retained. This practice differs from that used by Malikic et al. (2015), who selected the clone set containing genotypes closest to the true solution. Our approach is more practical because ground truth is not known in real world data analysis, which means that all equally likely solutions must be considered plausible.

We scored the number of correct clones (*n*_C_), i.e., those with *G*_E_ = 0. The proportion of true clones not correctly inferred is equal to %*E*_C_ = (1 – *n*_C_/N) × 100, where N was the number of true clones simulated. Some methods underestimated the number of clones. To negate this shortcoming, we computed the proportion of inferred clones that were not correct: %*E*_I_ =(1– [*n*_C_/*n*) × 100, where *n* is the number of clones inferred. We then computed the average of %*E*_C_ and %*E*_I_ (%*E*). All %*E*, %*E*_C_ and %*E*_I_ will be the same when the number of clones inferred is correct, i.e., *n* = N.

### 2.4 Existing inference methods evaluated and parameter settings

We applied all nine methods by using default and/or recommended settings and parameters (see Supplementary Note). The computation times required for some methods and datasets were extremely long (more than two weeks), prohibiting testing of all methods on all datasets. Also, some methods did not produce clone predictions for some of the datasets due to lack of convergence and other technical problems.

## 3 Results

### 3.1 Analysis of simulated datasets

We first present results from the analysis of G7 datasets in which the read counts were ascertained perfectly (i.e., no read count errors for any SNV) and the coverage was uniform for all the SNVs. This is an ideal situation in which methods are expected to perform the best. LICHeE and CloneFinder frequently produced the correct number of clone genotypes for these datasets (*n*; Table 2). Neither of these methods required users to specify the expected number of clones, unlike BayClone2, Canopy, and Clomial. For these, we input the correct number of clones, but they often produced multiple identical clone genotypes (with different clone names) and clones without any mutation assignment (normal cell). This resulted in the discovery of fewer than seven distinct clones. AncesTree and cloneHD also underestimated the number of clones. CITUP and Phy-loWGS overestimated the number of clones, because they produced multiple solutions (sets of clones) without any guidance to prefer one over the other. PhyloWGS produced five solutions (i.e., five sets of clones) per dataset, each containing 7 – 10 clones. CITUP produced six solutions per dataset, each containing eight clones.

**Table 2.** Performance for G7 datasets

| Method | $n$ | $n_c$ | Correct clones not found | Incorrect clones found | Average error |
| --- | --- | --- | --- | --- | --- |
| | | | $\%E_c$ | $\%E_i$ | $\%E$ |
| AncesTree | 5.0 | 2.0 | 71 | 60 | 66 |
| *BayClone2 | 6.8 | 3.5 | 50 | 48 | 49 |
| *Canopy | 6.3 | 2.8 | 60 | 56 | 58 |
| **CITUP | 15.0 | 7.0 | 0 | 53 | 27 |
| *Clomial | 6.8 | 3.7 | 47 | 46 | 47 |
| CloneFinder | 6.9 | 6.5 | 7 | 6 | 6 |
| cloneHD | 3.9 | 3.0 | 57 | 23 | 40 |
| LICHeE | 7.0 | 4.4 | 38 | 38 | 38 |
| **PhyloWGS | 8.8 | 4.0 | 42 | 54 | 48 |

\*True number of clones was specified in the input.
\*\*Multiple solutions (different sets of clones) can be produced for a dataset. All clones were pooled and redundant clones were eliminated.

We determined the error rate by scoring the number of true clones that were not found in the final collection of inferred clones (%*E*_C_). Most of the methods detected ~50% of the true clones (Table 2). For example, on average AncesTree inferred five clones, of which only two were identical to the correct clone genotypes (Table 2). CITUP and CloneFinder showed the strongest performance. For CITUP, %*E*_C_ was zero, i.e., all the correct clone genotypes were present in the final collection. But CITUP also discovered eight incorrect clones, which resulted in a large %*E*_I_ (Table 2). In contrast, CloneFinder showed the smallest %*E*_C_ and %*E*_I_. cloneHD also showed a relatively small %*E*_I_, but it only predicted a few clones, which resulted in a high %*E*_C_ (Table 2).

We allowed for 1, 2, and 3 SNV assignment errors when pairing inferred clones with true clone genotypes. This did not extensively change the clone count accuracy patterns (Fig. 4, solid lines), because the average clone genotype error was already fairly high for most of the methods, except CloneFinder (Fig. 5a). The highest level of genotype error (*G*_E_) was observed for AncesTree, and all other methods (excluding CloneFinder) showed similar performance (8% - 11% error) (Fig. 5a). Greater than 20% of the inferred clone genotypes produced by methods other than CloneFinder differed from the true clones by *G*_E_ > 10% (Fig. 5b).

**Figure 4:**
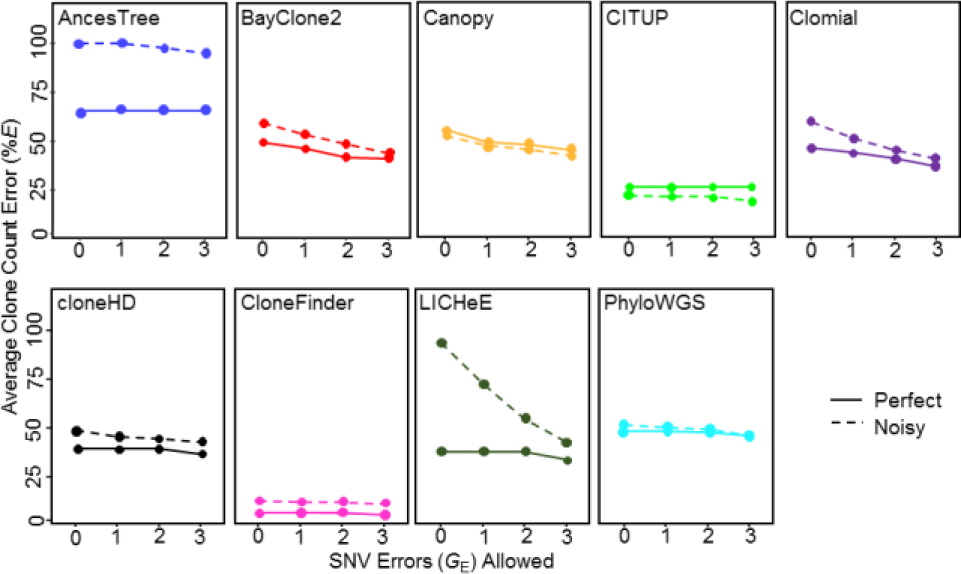
Average clone count errors in the analysis of G7 datasets. Average of the proportion of correct clones not found (%*E*_C_) and incorrect clones found (%*E*_I_), i.e., %*E*, were computed when allowing for one or more SNV assignment errors (*G_E_*). A solid line is for datasets without read count errors (perfect), and a dashed line is for datasets with read count errors (noisy).

We also examined the degree to which errors in SNV read counts (sampling errors) impacted method performance (Fig. 4, dashed line and Supplementary Table S1). Average of %*E*_C_ and %*E*_I_ became much higher for AncesTree when read counts were less accurate, but others performed similarly to the example analysis of perfect data, especially when we allowed a few genotype errors (Fig. 4). Most methods showed genotype errors similar to those for datasets without read counts error, except AncesTree which produced almost all clones with *G*_E_ > 5% (Fig. 5c and d). Overall, the average %*G*_E_ was less than 5% for CloneFinder and larger (~10%) for all other methods.

**Figure 5:**
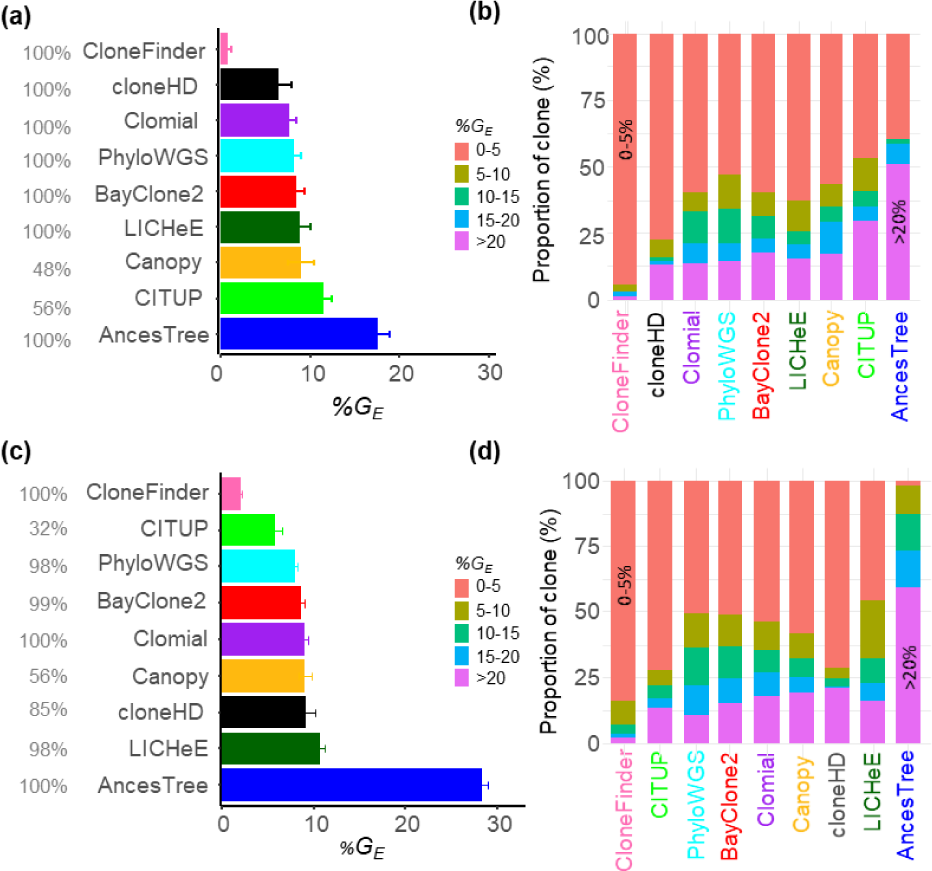
Performance of different methods for G7 datasets. Average clone genotype error with standard error for datasets in which SNV frequencies were known (**a**) without and (**c**) with read count errors. The proportion of datasets producing results are shown in gray. Proportion of clones in different %*G*_E_ categories for datasets in which SNV frequencies were known (**b**) without and (**d**) without read count errors. Methods are ordered from the best to worst (top to bottom and left to right).

We conducted simulation experiments where the number of SNVs was increased 10-fold (~1,000), consistent with whole exome/genome sequencing data. We changed the branch lengths of the 7-clone phylogenies (Fig. 1a) into 80 – 200 mutations (1,040 SNVs in total), i.e., new clones arose by accumulating 80 – 200 mutations. We found that LICHeE, CloneFinder, Clomial and cloneHD showed a performance similar to that observed for the G7 datasets (Supplementary Table S2). However, Phy-loWGS and AncesTree showed much lower accuracies, and BayClone2, Canopy, and CITUP suffered from technical problems and did not produce results.

Many contemporary datasets contain large numbers of samples, e.g., (Ross-Innes et al., 2015; Uchi et al., 2016), so we further tested our ability to analyze and discover correct clones using G12 datasets containing 11 tumor samples and 12 clones. Application of some methods was technically challenging for these datasets. For example, the maximum number of clones that CITUP and BayClone2 could infer was smaller than the true number of clones present. Clomial required extremely long computational times (> 2 week per dataset), prohibiting analyses of these larger datasets.

The analysis of G12 datasets produced results largely consistent with those seen for G7 datasets (Fig. 6a). The clone count error was the lowest for both CloneFinder and LICHeE, as the correct number of clones were produced (13 and 12 clones, on average, respectively) (Supplementary Table S3). Almost all clone genotypes inferred by CloneFinder and LICHeE had a few SNV assignment errors and, consequently, %*E* (average of %*E*_C_ and %*E*_I_) was very small (Fig. 6a). However, cloneHD produced only four clones on average and the average of %*E*_C_ and %*E*_I_ was very high. The clone count was always overestimated by PhyloWGS. Also, CloneFinder and LICHeE showed the highest accuracy, producing clone genotypes with 1% and 2% errors, respectively (Fig. 7). All other tested methods produced clones with considerably higher genotype error (8–19%) consistent with observations from the G7 datasets.

**Figure 6:**
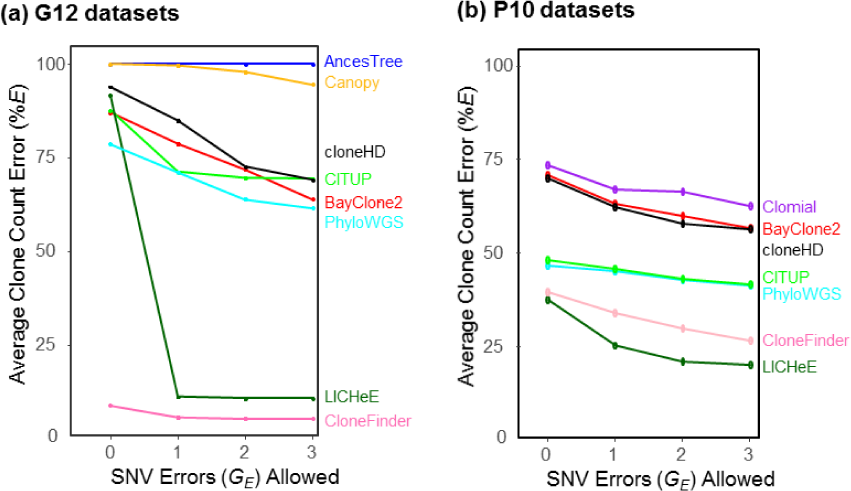
Average clone count errors in the analysis of G12 and P10 datasets. Average of the proportion of correct clones not found (%*E*_C_) and incorrect clones found (%*E*_I_) were computed when allowing for zero, one, two and three SNV assignment errors.

**Figure 7:**
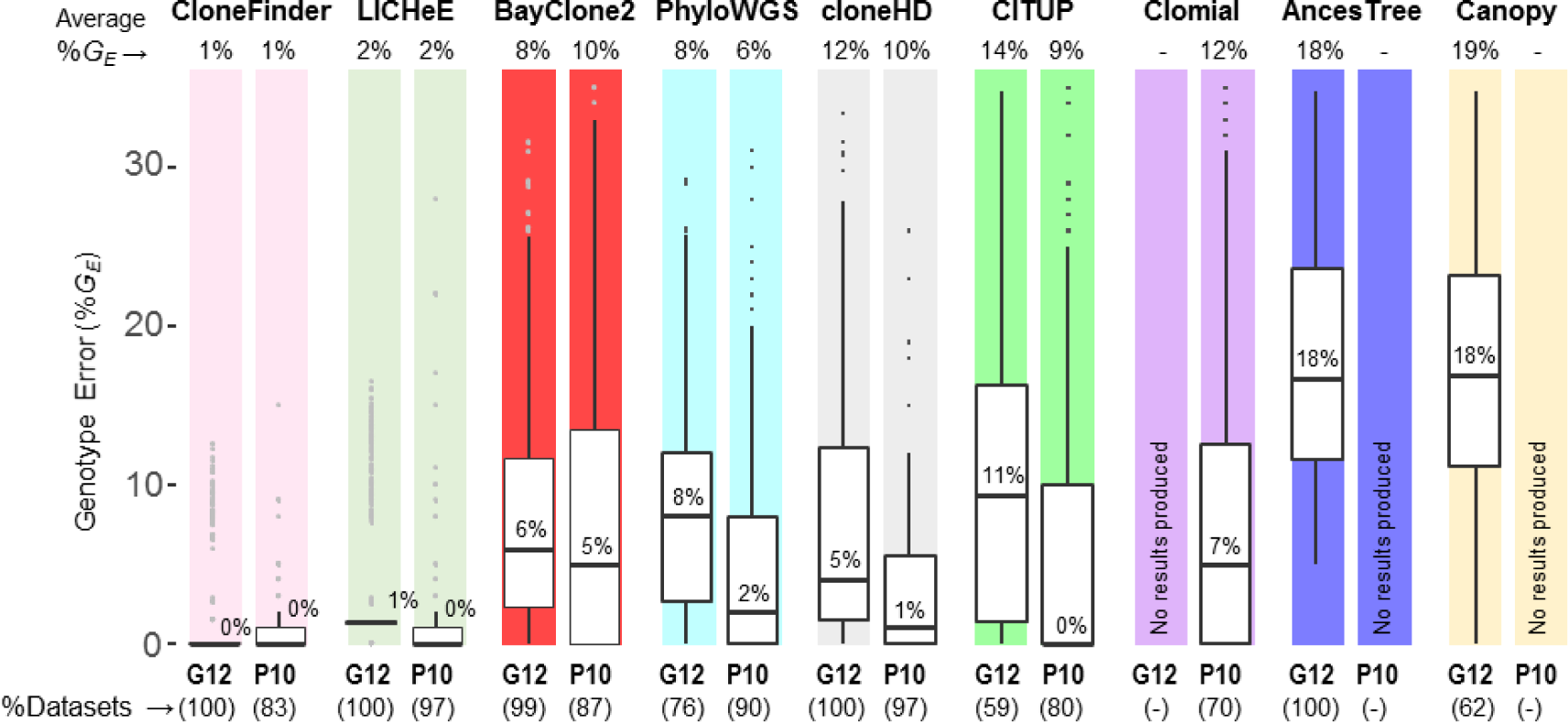
Performance of different methods for G12 and P10 datasets. Methods are ordered from best to worst (left to right), with *G*_E_ >35% falling outside the top range not shown. The average %*G*_E_ is given at the top and the median %*G*_E_ is shown within each boxplot. Percent datasets for which each method produced clones are shown at the bottom in the parentheses.

The above results are from datasets that were generated using only two clone phylogenies, so we next examined the performance of all the methods for the P10 datasets in which 10 different clone phylogenies were used (3 datasets each; see Fig. 3). Each P10 dataset contained six tumor samples, and the number of clones in each dataset varied from 6 to 10. As observed for G7 and G12 data, cloneHD regularly underestimated clone number (3 or 4 clones per dataset), whereas PhyloWGS could considerably overestimate the number of clones (up to 25 clones) (Supplementary Table S4). In contrast to results for G7 datasets, LICHeE and CloneFinder now underestimated clone counts, but still produced the smallest %*E* among the nine methods tested (Fig. 6b).

In terms of genotype error, the analysis of P10 datasets showed patterns similar to those observed for G12 data (Fig. 7). The average %*G*_E_ of clones produced by LICHeE and CloneFinder was small (<2%), which means that the clones produced by these two methods were generally accurate even though they did not always discover all the clones. Similarly, the average %*G*_E_ of PhyloWGS was low (6%). Although the average %*G*_E_ of CITUP and cloneHD was higher, the median of %*G*_E_ was low (0% and 1%, respectively). Thus, these methods produced many clone genotypes accurately, but a few clone genotypes contained high error. The other methods produced *G*_E_ in the range of 9 – 12%, as around quarter of inferred clones had *G*_E_ >15%, consistent with observations from the G7 and G12 analyses.

We next tested E10 datasets, where the clone sampling scheme differed from P10 datasets. In E10 datasets, tumor samples were populated with 4 – 6 clones that were randomly selected from the clone phylogeny (El-Kebir et al., 2015). In contrast to P10 datasets, this strategy allows the coexistence of any clones within a sample, and such a situation may arise when cancer cells often migrate to another section of a tumor or between tumors. Popic et al. (2015) refer to this higher level of clone mixing in tumor samples to be “randomized sampling.”

We begin with an analysis of the performance of AncesTree, because the E10 datasets were originally used to test the AncesTree method (El-Kebir et al., 2015). Surprisingly, AncesTree did not perform well in predicting the number of clones and inferring clone genotypes (Supplementary Table S5 and Fig. 8). Also, it performed worse than CITUP and PhyloWGS, which conflicts with the report (El-Kebir et al., 2015) that AncesTree’s performance was similar to or better than CITUP and PhyloSub (the precursor of PhyloWGS). This difference is partly explained by the fact that the original study did not consider the large number of unassigned SNVs (lacking assignment to any inferred clones; median of 31 of the 100 total variants) when measuring accuracy. This treatment is likely to have overestimated the accuracy of AncesTree in the original study. We assigned wild-type alleles to all these sites, as there was no justification for imposing variants.

**Figure 8:**
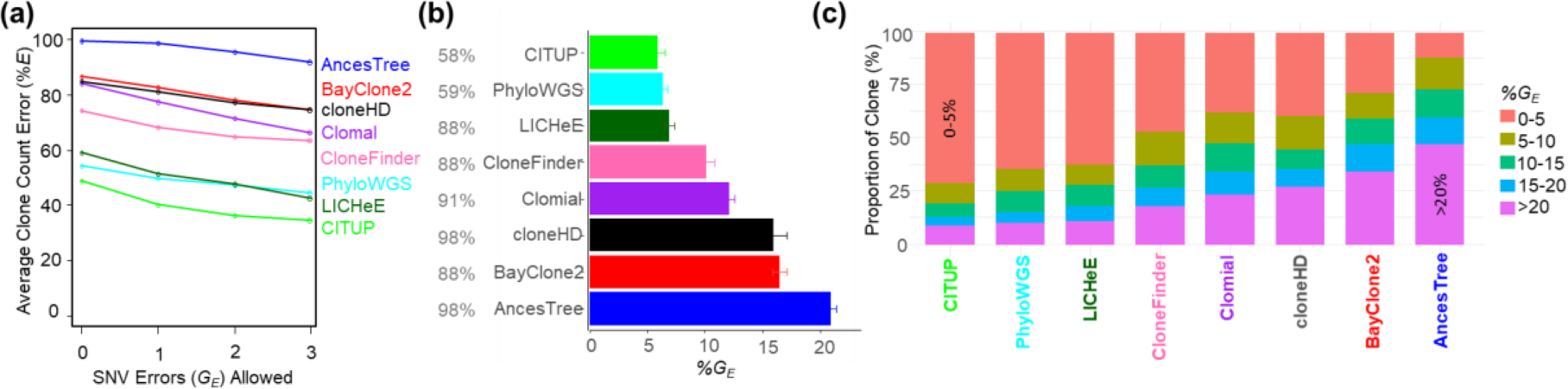
Performance of clone prediction methods for E10 datasets. (**a**) Average error in the inference of the number of clones when allowing up to 3 SNV assignment errors. (**b**) Average clone genotype error, along with their standard errors. Methods are ordered from best to worst. (**c**) Proportion of clones with various degrees of genotype errors. We used 90 datasets, and the number of datasets analyzed by each method is shown in gray before the method name in panel b. Canopy failed to analyze all the datasets.

No method performed well for these datasets, as the average of %*E*_C_ and %*E*_I_ was rather high (Fig. 8a). Even the best performing methods (CITUP, LICHeE, and PhyloWGS) did not achieve an average %*G*_E_ less than 5% (Fig. 8b) and around a quarter of inferred clones showed *G*_E_ > 10% (Fig. 8c). Therefore, clone prediction methods do not perform well when the tumor samples contain random mixtures of clones.

### 3.2 Analysis of empirical datasets

We analyzed two empirical datasets. The first empirical dataset was obtained from Stachler et al. (2015). It consisted of two samples from esophageal adenocarcinoma, five samples from precursor lesions, and one metastatic sample from the lymph node (Patient 7). This dataset contained 683 SNVs with estimated CCFs. Although the average total read count was relatively high (88) in this dataset, many SNV positions had a low read coverage (<50 total reads per sample). Such positions suffer from false-negative and false-positive detection of mutations (Reiter et al., 2017). Indeed, many SNVs with <50 read count did not obtain strong support for the presence/absence of mutations when analyzed by Reiter et al. (2017)’s method, as the probability of presence of mutations was less than 95%. Therefore, we removed all such SNVs if they were supported by fewer than 50 reads in more than half of the tumor samples. This reduced the dataset to 430 SNVs.

For this dataset, CloneFinder inferred that one sample of low-grade dysplasia contained the root clone, which was consistent with the original suggestion by Stachler et al. (2015). However, CloneFinder produced 11 clones (Fig. 9a). This count is smaller than the number of clones reported in the original study (16 clones) because CloneFinder failed to infer a few tumor-specific clone genotypes reported in Stachler et al. (2015). This is consistent with our observations for simulated E10 datasets, in which CloneFinder had difficulty deconvoluting clone genotypes. Among other methods, cloneHD greatly underestimated the number of clones (four clones), consistent with computer simulation results. PhyloWGS produced five solutions, each containing a large number of clones (42 - 51 clones), as observed in our analyses of simulated data. LICHeE, Canopy and AncesTree failed to produce any results; BayClone2 did not converge with 10,000 samples; Clomial and CITUP did not finish computation even after two weeks. Thus, only three of nine methods evaluated were capable of completing analysis of this data. In addition, the original study (Stachler et al. (2015)) did not report clone genotypes, precluding comparison of the clone genotypes produced by the three completed methods to the original study.

**Figure 9:**
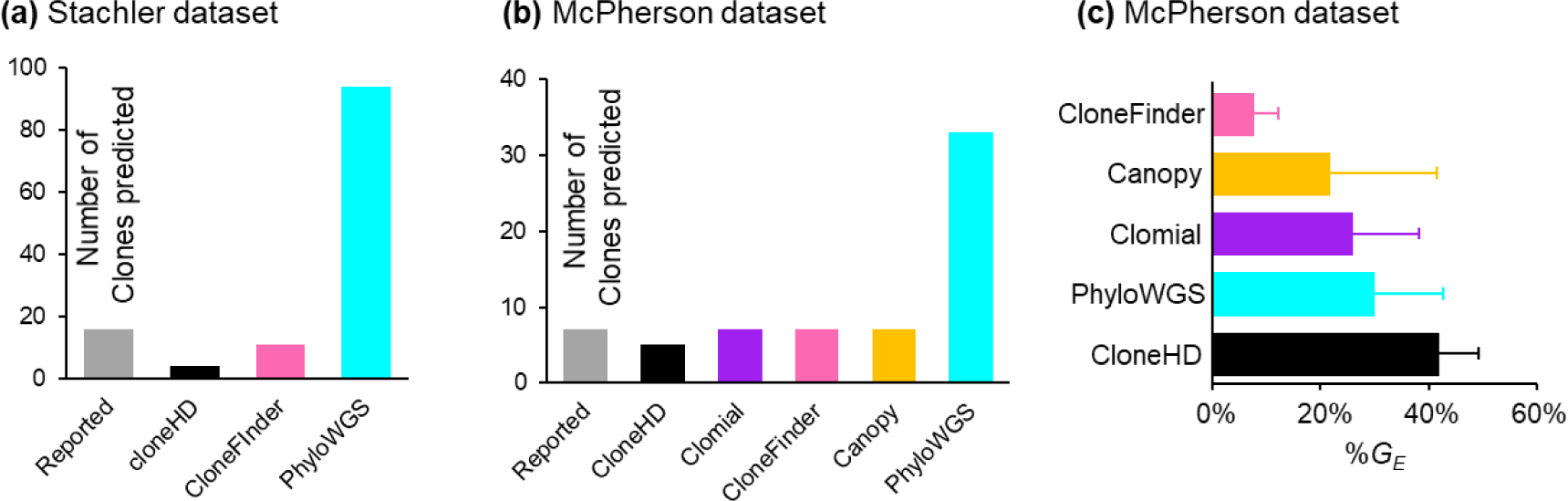
Results from the analysis of two empirical datasets. Number of clones found by different method for (**a**) Stachler et al. (2015) dataset and (**b**) McPherson et al. (2016) dataset. (**c**) Average clone genotype difference (%*G*_E_) between the clone genotypes inferred by McPherson et al. (2016) and those produced by different methods. Standard deviation bars quantify the degree of dispersion in *G*_E_.

We next discuss results from the analysis of a dataset from McPherson et al. (2016), which contained 130 SNVs for 11 tumor regions (samples) of a high-grade serous ovarian cancer (patient 3). The authors identified seven clone genotypes from bulk-sequencing data and validated them using targeted single-cell sequencing. We inferred clone genotypes from their bulk sequencing data, and then paired each of the inferred clone genotype with the most similar clone genotype reported in McPherson et al. (2016). In this comparison, we excluded all SNVs (38%) that were predicted to be affected by loss/recurrence of mutations due to CNAs by McPherson et al. (2016).

We evaluated whether the nine tested clone prediction methods produced clone genotypes similar to those validated by single-cell sequencing results in the original study of McPherson et al. (2016). For this dataset, CloneFinder inferred seven clones, which agreed with the original study (McPherson et al., 2016) (Fig. 9b). The average *G*_E_ was 7%, which was the lowest of all methods tested (Fig. 9c). Canopy produced seven clones with an average *G*_E_ equal to 22%, but three clone genotypes showed *G*_E_ < 10%. None of the seven clones produced by Clomial showed *G*_E_ < 10%, and all clones produced by cloneHD (5 clones) showed *G*_E_ > 32%. PhyloWGS produced five solutions, each containing 30 clones, which was consistent with our conclusion based on results from simulated datasets: PhyloWGS often produces too many clones. No results from LICHeE, AncesTree, BayClone2 and CITUP were available, because they all failed at various stages of computation. While the above tests of the performance of clone prediction methods by using empirical datasets is useful, we believe that targeted studies, where the accuracy of the inferred clone genotypes is directly tested via single-cell sequencing, are necessary to further evaluate methods. This is because base assignment in single-cell sequencing is not very accurate at present (Gawad et al., 2016; Navin, 2015; Sun et al., 2015) and that some clones may not be sampled when only a few cells are singly sequenced.

## 4 Discussion

CloneFinder and LICHeE performed best when individual tumor samples consisted of clones from the same evolutionary lineage (e.g., G7, G12, and P10 datasets). In CloneFinder, this property leads to informative initial tumor phylogenies and allow reliable inference of root and ancestral clone genotypes. LICHeE also uses the patterns of mutation presence and absence among tumor samples to generate SNV clusters and thus harnesses evolutionary information in the data. We expect many real world datasets to exhibit this evolutionary property in which similar clones are inherited by descendant tumors and distantly-related clones are found in samples from distantly located tumor or regions of a tumor, which means that tumor expansion is coupled with the evolution of new clones (Davis et al., 2017; Gerlinger et al., 2014; Gerlinger et al., 2012; Hu et al., 2017). Many other methods did not perform well for these datasets, because they seem to not use the intrinsic evolutionary information effectively.

Across methods, the accuracy of clone counts and genotypes was much lower for datasets in which all tumor samples contained random mixtures of clones. Such data would be produced for liquid tumors and solid tumors with disordered tumor growth (Sottoriva et al., 2015), or if tumor cells frequently migrate within or between tumors (E10 datasets). In this case, the relatedness of clones within a tumor sample will not be significantly greater than that between samples. For such datasets, CloneFinder did not perform well. However, the average %*G*E was relatively high (>5%) even for the best methods, suggesting that there is still a lot of room for improvement in all methods developed to date.

At the same time, the performance of all the methods requires further evaluation. The accuracy of clone predictions will be influenced by many additional factors not simulated for this study, including the evolution of tumors and clones under strong natural selection, the presence of CNAs, sequencing errors, effects of different read depths, and mutational homoplasy (backward, parallel, and multiple hits at a position). In addition, there is an urgent need for improved software implementations of current methods, because many methods failed to produce results for a large number of datasets.

Direct comparison of nine methods used to analyze the same collection of simulated datasets revealed that many current methods did not perform as well as previously reported. Therefore, we discuss similarities and differences of our results with prior reports. To begin with, LICHeE performed well on G12 and P10 data in our simulation study, but its accuracy was limited on E10 data. Popic et al. (2015) also generated datasets that were similar to G12/P10 and E10, which they called localized and randomized sampling, respectively. Similar accuracies were reported for both types of datasets, but accuracy was evaluated by examining the relative order of mutations, and did not include clone genotype errors. Our tests of LICHeE provide a useful benchmark of its absolute and relative performance, as currently LICHeE has only been rigorously compared in peer-reviewed literature with Treeomics, a software which does not infer clone genotypes (Reiter et al., 2017).

Jiang et al. (2016) reported that the proportion of SNV assignment errors per dataset was generally less than 10% for Canopy, a result that is consistent with our observations here. Canopy never produced perfect results for any dataset in our analyses, but Jiang et al. (2016) indicated that Canopy could generate all clones perfectly for many simulated datasets. We suspect that this difference reflects the application of Canopy to smaller datasets (three clones in three tumor samples). Jiang and coauthors also suggested that the clone genotype error rates become smaller (< 4%) when the number of SNVs were larger (up to 200), but we found Canopy to have *G*_E_ >5%. The higher error rate that we observed could be caused by different parameter settings. Canopy also failed to produce results for a large fraction of datasets. These results may reflect a strong sensitivity of this method to selection of parameter settings, where the consequence of a poor choice is complete failure.

PhyloSub (Jiao et al., 2014), the predecessor of PhyloWGS, has been subject to frequent comparison with other methods, so we use its accuracy as a proxy for PhyloWGS. For example, PhyloSub was compared with CITUP by Malikic et al. (2015), who reported superior performance of CITUP. However, our simulation studies did not show much difference between these methods, which is likely due to the treatment of multiple equally likely results. Among equally likely solutions, Malikic et al. (2015) selected the solution that showed the best accuracy. Indeed, we also observed one perfect solution from CITUP for our 7-clone simulation (without read count errors). However, as mentioned earlier, there is no practical way to select the correct result from all the equally likely results when analyzing empirical data, and thus the evaluation of CITUP in the previous study did not reflect realistic performance.

Zare et al. (2014) reported that all clones predicted by Clomial were nearly perfect for datasets with a small number of clones (3 clones) and a larger number of samples (>5 samples). However, Clomial did not show high accuracy for most of our datasets, probably due to the larger number of clones relative to the number of samples tested. In fact, when the number of clones was larger, Zare et al. (2014) also reported lower clone genotype accuracy (~10% error rates), similar to our findings. Jiang et al. (2016) also reported similar error rates for Clomial (5 – 7%). It is, however, important to note that the accuracy of Clomial can be improved by selecting the most likely model after evaluating a very large number of models by using many different initial values. In fact, the average %*G*_E_ across inferred clones for a 7-sample simulation with read count errors improved dramatically with 1,000 models (decreasing to 4% from 9–11%). However, in practice, the number of models that is necessary is unknown, and the use of many models is computationally exhausting (~3 weeks for this relatively small 7-sample simulation with 1,000 models). Whenever possible, it is best to run Clomial with a very large number of initializations. As for cloneHD and BayClone2, the original articles employed small datasets (only two clones), while our analyses were performed on datasets containing up to 12 clones. BayClone2 also simulated four clones, but the number of samples was very large (25 samples). Therefore, these methods may perform better for such special cases where the number of samples far exceeds the number of clones. Our results provide further insights into cloneHD and BayClone2 performance for more realistically-sized datasets.

In summary, current methods to infer clone genotypes from SNV allele frequencies, produced by using bulk sequencing techniques, show moderate accuracies for simulated datasets. This information will be useful when interpreting results from the application of these computational tools, and it highlights areas in need of improvement.

## Acknowledgements

We thank Drs. Jeffrey Townsend, Camille Ragin, Zhai Weiwei, Glen Stecher, Craig Stanley and Adam Khalili for critical comments and technical support.

## Funding

Grants from Temple University and from the National Institutes of Health to S.K. (LM012487) and S.M. (LM012758) provided support for this research.

*Conflict of Interest:* none declared.

